# Enhancing bioreactor arrays for automated measurements and reactive control with ReacSight

**DOI:** 10.1101/2020.12.27.424467

**Authors:** François Bertaux, Sebastián Sosa-Carrillo, Achille Fraisse, Chetan Aditya, Mariela Furstenheim, Gregory Batt

## Abstract

New small-scale, low-cost bioreactors provide researchers with exquisite control of environmental parameters of microbial cultures over long durations, allowing them to perform sophisticated, high-quality quantitative experiments that are particularly useful in systems biology, synthetic biology and bioengineering. However, existing setups are limited in their automated measurement capabilities, primarily because sensitive and specific measurements require bulky, expensive, stand-alone instruments. Here, we present ReacSight, a generic and flexible strategy to enhance bioreactor arrays for automated measurements and reactive experiment control. On the hardware side, ReacSight leverages a pipetting robot for sample collection, handling and loading. On the software side, ReacSight provides a versatile instrument control architecture and a generic event system for reactive experiment control. ReacSight is ideally suited to integrate open-source, open-hardware components but can also accommodate closed-source, GUI-only components (e.g. cytometers). We use ReacSight to assemble a platform for cytometry-based characterization and reactive optogenetic control of parallel yeast continuous cultures. Using a dedicated bioreactor array, we showcase its capabilities on three applications. First, we achieve parallel real-time control of gene expression with light in different bioreactors. Second, we explore the impact of nutrient scarcity on fitness and cellular stress using well-controlled, high-information content competition assays. Third, we exploit nutrient scarcity to achieve dynamic control over the composition of a two-strain consortium. To illustrate the genericity of ReacSight, we also assemble an equivalent platform using the optogenetic-ready, open-hardware and commercially available Chi.Bio bioreactors.

## Introduction

Small-scale, low-cost bioreactors are emerging as powerful tools for microbial systems and synthetic biology research^1–4^. They allow tight control of cell culture parameters (e.g. temperature, cell density, media renewal rate) over long durations (several days). These unique features enable researchers to perform sophisticated experiments and to achieve high experimental reproducibility. Examples include characterization of antibiotic resistance when drug selection pressures increases as resistance evolves^1^, cell-density controlled characterization of cell-cell communication synthetic circuits^2^, and genome-wide characterization of yeast fitness under dynamically changing temperature using a pooled knockout library^3^.

A weakness of existing small-scale, low-cost bioreactors is their limited automated measurement capabilities: in situ optical density measurements only inform about overall biomass concentration and its growth rate, and, when available^2,4^, fluorescence measurements suffer from low sensitivity and high background. It is often essential to also measure and follow over time key characteristics of the cultured cell population, such as gene expression levels, cellular stress levels, cell size and morphology, cell cycle progression, proportions of different genotypes or phenotypes. Researchers usually need to manually extract, process and measure culture samples to run them through more sensitive and specialized instruments (e.g. a cytometer, a microscope, a sequencer). Manual interventions are usually tedious, error-prone and strongly constrains the available temporal resolution and scope (i.e. no time-points during night-time). It also impedes the dynamic adaptation of culture conditions in response to such measurements. Such *reactive experiment control* is currently gaining interest in systems and synthetic biology. It can be used to either maintain a certain state of the population (*external feedback control*) or to maximize the value of the experiment (*reactive experiment design*). For example, external feedback control can be used to disentangle complex cellular couplings and signaling pathway regulations^5–8^, to steer the composition of microbial consortia^9^ or to optimize industrial bioproduction^10^. Reactive experiment design can be especially useful in the context of long and uncertain experiments such as artificial evolution experiments^11^. It is also useful to accelerate model-based characterization of biological systems by enabling real-time parameter inference and optimal experiment design^12^.

In principle, commercial robotic equipment and/or custom hardware can be used to couple a bioreactor array to a sensitive, multi-sample (typically accepting 96-well plates as input) measurement device. However, this poses tremendous challenges regarding equipment sourcing, equipment cost and software integration. When a functional platform is established, upgrade and maintenance of the corresponding hardware and software is also highly challenging. Accordingly, very few examples have been reported to date. For instance, only two groups have demonstrated automated cytometry and reactive optogenetic control of bacteria^13^ or yeast^6,7^ cultures, with setups limited to either a single continuous culture or multiple batch-only cultures. One group has also demonstrated automated microscopy and reactive optogenetic control of a single yeast continuous culture^14^.

Here, we present ReacSight, a generic and flexible strategy to enhance bioreactor arrays for automated measurements and reactive experiment control. We first use ReacSight to assemble a platform enabling cytometry-based characterization and reactive optogenetic control of parallel yeast continuous cultures. Importantly, we built two versions of the platform, using either a custom-made bioreactor array or the recent low-cost, open-hardware, optogenetic-ready commercially available Chi.Bio bioreactors^4^. We then demonstrate its usefulness on three case studies. First, we achieve parallel real-time control of gene expression with light in different bioreactors. Second, we explore the impact of nutrient scarcity on fitness and cellular stress using highly-controlled and informative competition assays. Third, we exploit nutrient scarcity and the reactive experiment control capabilities of the platform to achieve dynamic control over the composition of a two-strain consortium. To the best of our knowledge, this last application is the first of its kind.

## Results

### Measurement automation, platform software integration and reactive experiment control with ReacSight

The ReacSight strategy (Figure 1, Text S1.1) to enhance bioreactor arrays for automated measurements and reactive experiment control combines hardware and software elements in a flexible and standardized manner. A pipetting robot is used to establish, in a generic fashion, a physical link between any bioreactor array and any plate-based measurement device (Figure 1, left). Bioreactor culture samples are sent to the pipetting robot through pump-controlled sampling lines attached to the robot arm (*sampling*). A key advantage of using a pipetting robot is that diverse treatment steps can be automatically performed on culture samples before measurement (*treatment)*. Samples are then transferred to the measurement device by the pipetting robot (*loading*). Naturally, this requires that the measurement device can be physically positioned such that when its loading tray is open, wells of the device input plate are accessible to the robot arm. Partial access to the device input plate is not problematic because the robot can be used to wash input plate wells between measurements, allowing re-use of the same wells over time (*washing*). Importantly, if reactive experiment control is not needed or if it is not based on measurements, the robot capabilities can also be used to treat and store culture samples for one-shot offline measurements at the end of an experiment, enabling automated measurements with flexible temporal resolution and scope.

**Figure 1.**
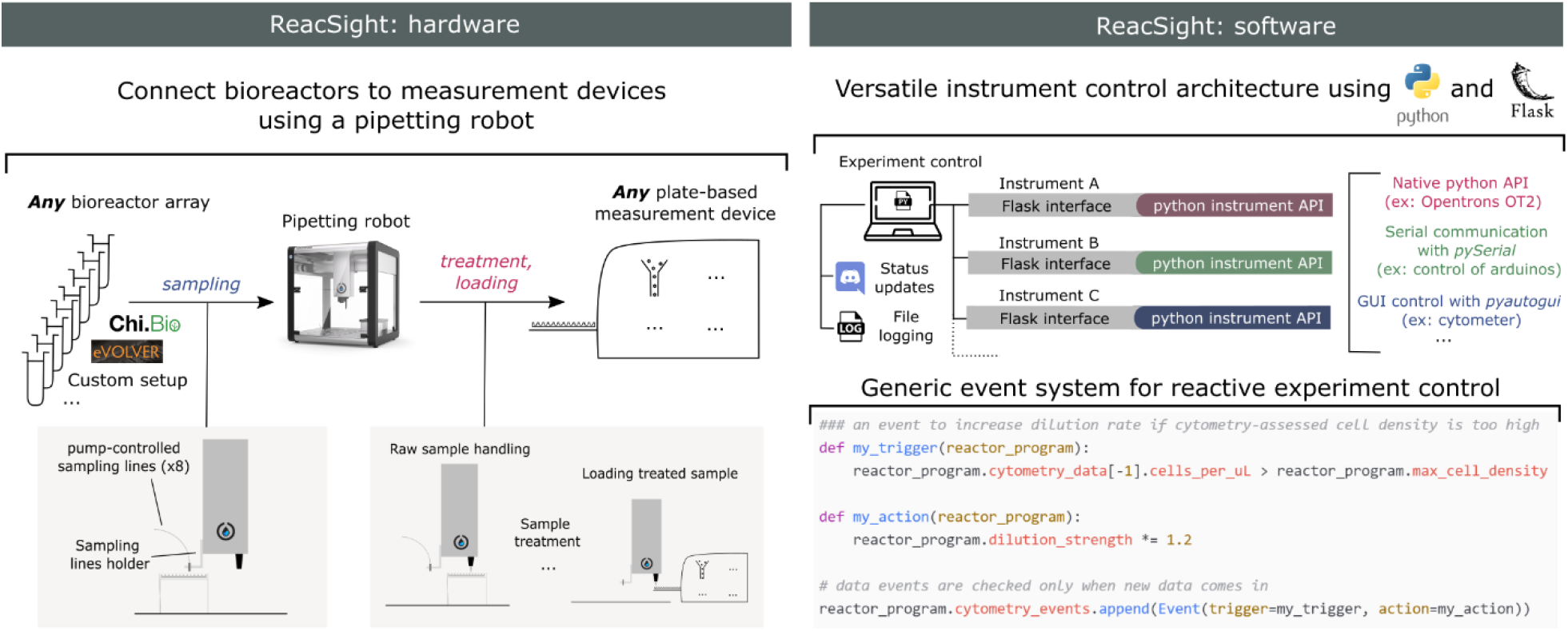
ReacSight: a strategy to enhance bioreactor arrays for automated measurements and reactive experiment control. On the hardware side, ReacSight leverages a pipetting robot (such as the low-cost, open-source Opentrons OT-2) to create a physical link between any multi-bioreactor setup (eVOLVER, Chi.Bio, custom…) and the input of any plate-based measurement device (plate reader, cytometer, high-throughput microscope, pH-meter…). If necessary, the pipetting robot can be used to perform a treatment on bioreactor samples (dilution, fixation, extraction, purification…) before loading into the measurement device. If reactive experiment control is not needed, treated samples can also be stored on the robot-deck for offline measurements (the OT-2 temperature module can help the conservation of temperature-sensitive samples). On the software side, ReacSight enables full platform integration via a versatile instrument control architecture based on Python and the Python web application framework Flask. ReacSight software also provides a generic event system to enable reactive experiment control. Example code for a simple use case of reactive experiment control is shown. Experiment control can also inform remote users about the status of the experiment using Discord webhooks and generates an exhaustive log file.

ReacSight also provide a solution to several software challenges that should be addressed to unlock automated measurements and reactive experiment control of multi-bioreactors (Figure 1, right). First, programmatic control of all instruments of the platform (bioreactors, pipetting robot, measurement device) is required. Second, a single computer should communicate with all instruments to orchestrate the whole experiment. ReacSight combines the versatility and power of the python programming language with the genericity and scalability of the Flask web-application framework to address both challenges. Indeed, Python is ideally suited to easily build APIs to control various instruments: there exist well-established, open-source libraries for the control of micro-controllers (such as arduinos), and even for the ‘clicking’-based control of GUI-only software driving closed-source instruments lacking APIs (pyautogui). Importantly, the open-source, low-cost pipetting robot OT-2 (Opentrons) is shipped with a native Python API. Hamilton robots can also be controlled with a Python API^15^. Flask can then be used to expose all instrument APIs for simple access over the local network. The task of orchestrating the control of multiple instruments from a single computer is then essentially reduced to the simple task of sending HTTP requests, for example using the Python module requests. HTTP requests also enable user-friendly communication from the experiment to remote users using the community-level digital distribution platform Discord. This versatile instrument control architecture is a key component of ReacSight. Two other key components of ReacSight are 1) a generic object-oriented implementation of *events* (if *this* happens, do *this*) to facilitate reactive experiment control and 2) an exhaustive logging of all instrument operations into a single log file. ReacSight software as well as source files for hardware pieces are made openly available in the ReacSight Git [repository](https://gitlab.inria.fr/InBio/Public/reacsight).

### Reactive optogenetic control and single-cell resolved characterization of yeast continuous cultures

Our first application of the ReacSight strategy is motivated by yeast synthetic biology applications. In this context, it is critical to 1) accurately control synthetic circuits and 2) measure their output in well-defined environmental conditions and with sufficient temporal resolution and scope. Optogenetics provides an ideal way to control synthetic circuits, and bioreactor-enabled continuous cultures are ideal to exert tight control over environmental conditions for long durations. To measure circuit output in single cells, cytometry is also ideal due to high sensitivity and throughput. We thus resorted to the ReacSight strategy to assemble a fully automated experimental platform enabling reactive optogenetic control and single-cell resolved characterization of yeast continuous cultures (Figure 2A), using a benchtop cytometer as a measurement device.

**Figure 2.**
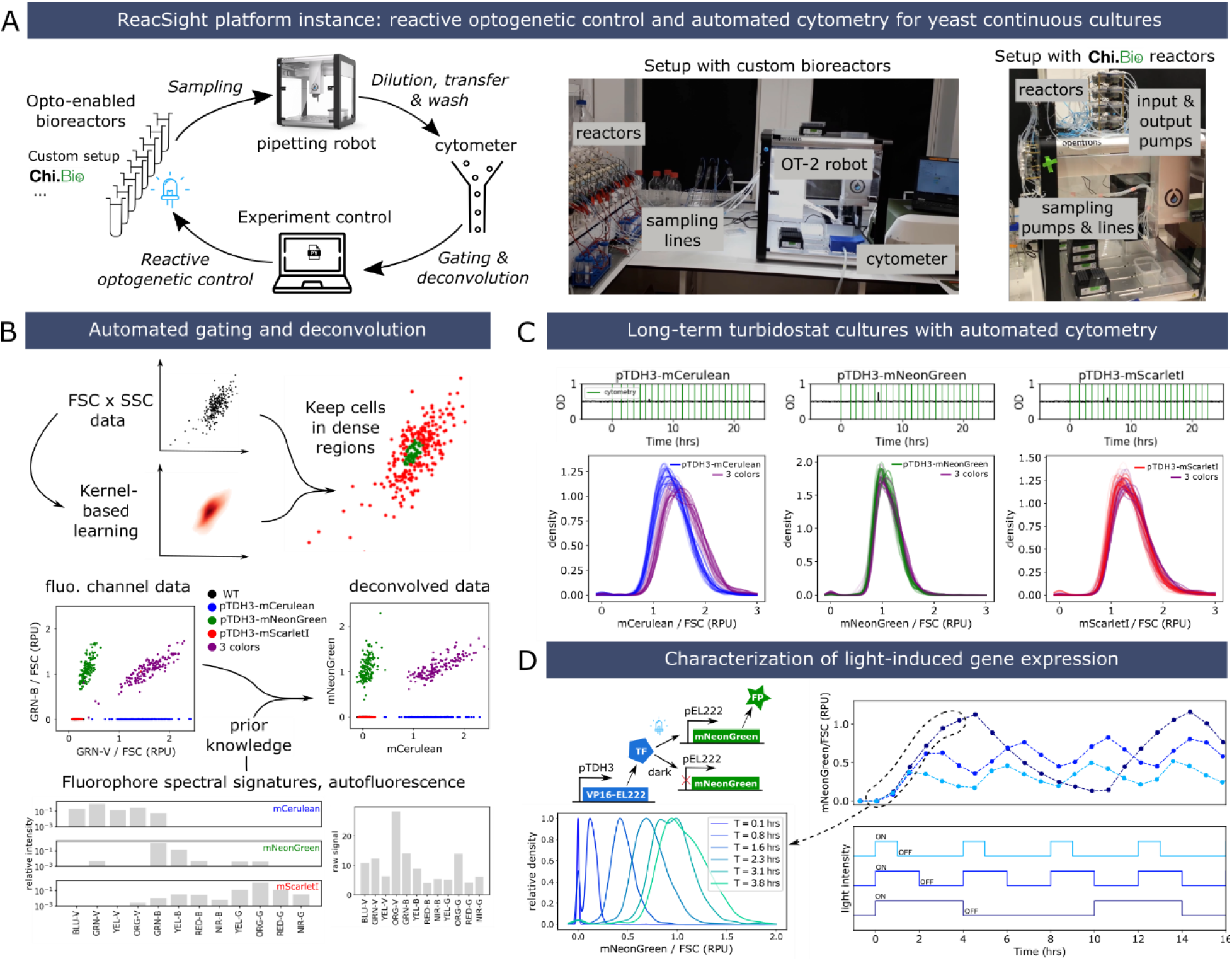
ReacSight-based assembly of a fully automated platform enabling reactive optogenetic control and single-cell resolved characterization of yeast continuous cultures. (A) Platform overview. The Opentrons OT-2 pipetting robot is used to connect optogenetic-ready multi-bioreactors to a benchtop cytometer (Guava EasyCyte 14HT, Luminex). The robot is used to dilute fresh culture samples in the cytometer input plate and to wash it between timepoints. The ‘clicking’ python library pyautogui is used to create the cytometer instrument control API. Custom algorithms were developed and implemented in python to automatically gate and deconvolve cytometry data on the fly. Two versions of the platform were assembled, using either a custom bioreactor setup (left photos) or Chi.Bio reactors4 (right photo). (B) Description of the gating and deconvolution algorithm. As an example, deconvolution between the overlapping fluorophores mCerulean and mNeonGreen are shown. (C) Stability of single-cell gene expression distributions over many generations. Strains constitutively expressing either mCerulean, mNeonGreen or mScarlet-I alone or altogether (‘3-colors’ strain) from the transcriptional units driven by the pTDH3 promoter and integrated in the chromosome were grown in turbidostat mode (OD setpoint = 0.5, upper plots) and cytometry was acquired hourly (vertical green lines). Distributions (smoothed via Gaussian kernel density estimation) of fluorophore levels (after gating, deconvolution, and normalization by the forward scatter, FSC) for all timepoints are plotted together with different color shades (bottom). RPU: relative promoter units (see Methods). (D) Characterization of a light-driven gene expression circuit based on the EL222 system16. Three different ON-OFF blue light temporal profiles were applied (bottom) and cytometry was acquired every 45 minutes. The median of gated, deconvolved, FSC-normalized data is shown (top). All bioreactor experiments presented in this figure were performed in parallel, the same day, with the custom bioreactor platform version.

Detailed information on the platform hardware and software is provided in Text S1.2, and we discuss here only key elements. Eight reactors are connected to the pipetting robot, meaning that each timepoint fills one row of a sampling plate. While three rows of the cytometer input plate are accessible by the robot, we use only one row, washed extensively by the robot to achieve less than 0.2% carry-over as validated using beads. We typically fit two tip boxes and two sampling plates (2 × 96 = 192 samples) on the robot deck, therefore enabling 24 timepoints for each of the 8 reactors without any human intervention. To enable reactive experiment control based on cytometry data, we developed and implemented algorithms to perform automated gating and spectral deconvolution between overlapping fluorophores (Figure 2B).

We first validated the performance of the platform by carrying out long-term turbidostat cultures of yeast strains constitutively expressing various fluorescent proteins from chromosomally integrated transcriptional units (Figure 2C). Distributions of fluorophore levels were unimodal and stable over time, as expected from steady growth conditions with a constitutive promoter. Distributions of mNeonGreen and mScarlet-I exactly overlapped between the single- and 3-color strains, as expected from the assumptions that expressing one or three fluorescent proteins from the strong pTDH3 promoter has negligible impact on cell physiology and that the relative positioning of transcriptional units in the 3-color strain (mCerulean first, followed by mNeonGreen and mScarlet-I) has little impact on gene expression. Measured levels of mCerulean appear slightly higher (~15%) in the 3-color strain compared to the single-color strain. This could be caused by residual errors in the deconvolution, exacerbated by the low brightness of mCerulean compared to autofluorescence and to mNeonGreen.

Finally, to validate the optogenetic capabilities of the platform, we built and characterized a light-inducible gene expression circuit based on the EL222 system^16^ (Figure 2D). As expected, applying different ON-OFF temporal patterns of blue light resulted in dynamic profiles of fluorophore levels covering a wide range, from near-zero levels (i.e., hardly distinguishable from auto-fluorescence) to levels exceeding those obtained with the strong constitutive promoter pTDH3. Cell-to-cell variability in expression levels at high induction is also low, with coefficient of variation (CV) values comparable to the pTDH3 promoter (0.22 vs 0.20).

The first platform we assembled used a pre-existing, custom optogenetic-enabled bioreactor array (Supplementary Text S1.2.1). This setup has several advantages (reliability, wide range of working volumes) but cannot be replicated easily by other labs. Thanks to the modularity of the ReacSight architecture, we could quickly construct a second version of the platform with similar capabilities by exchanging this custom bioreactor array with an array of the recently described, open-hardware, optogenetic-ready, commercially available Chi.Bio^4^ bioreactors (Figure 2A, right photo, Supplementary Text S1.2.2). To validate the performance of this other version of the platform, we performed optogenetic induction experiments with the same strain as in Figure 2D and obtained excellent reactor-to-reactor reproducibility for various light induction profiles (Figure 6B in Supplementary Text S1).

### Real-time control of gene expression using light

To showcase the reactive optogenetic control capabilities of the platform, we set out to dynamically adapt light stimulation so as to maintain fluorophore levels at different target setpoints. Such in-silico feedback for in-vivo regulation of gene expression is useful to dissect the functioning of endogenous circuits in the presence of complex cellular regulations and could facilitate the use of synthetic systems for biotechnological applications^6,10,17^.

We first constructed and validated a simple mathematical model of light-induced gene expression (Figure 3A). Joint fitting of the three model parameters to the characterization data of Figure 2D resulted in an excellent quantitative agreement. This is remarkable given the simplicity of the model assumptions: constant rate of mRNA production under light activation, constant translation rate per mRNA, and first-order decay for mRNA (mainly degradation, half-life of 20 minutes) and protein (mostly dilution, half-life of 1.46 hours). Therefore, when experimental conditions are well-controlled and the data is properly processed, one can hope to quantitatively explain the behavior of biological systems with a small set of simple processes. We then incorporated the fitted model into a model-predictive control algorithm (Figure 3B). Together with the ReacSight event system, this algorithm enabled accurate real-time control of fluorophore levels to different targets in different reactors in parallel (Figure 3C).

**Figure 3.**
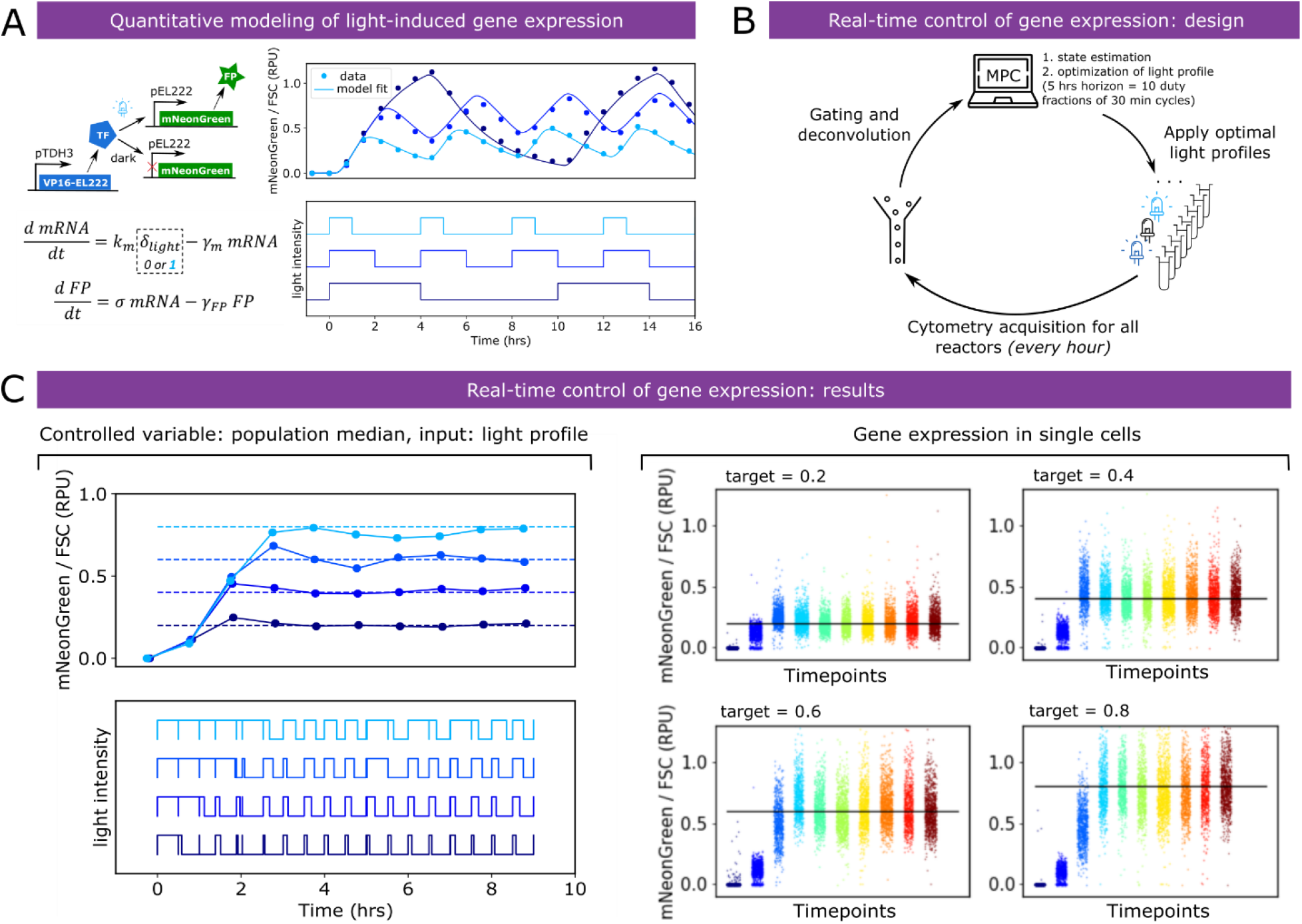
Closing the loop: real-time control of gene expression using light. (A) A simple ODE model of the light-driven gene expression circuit is fitted to the characterization data of Figure 2D. Fitted parameters are γ_m_ = 2.09 hr^−1^, σ = 0.64 RPU. hr^−1^ *and* γ_FP_ = 0.475 hr^−1^. k_m_ *was arbitrarily set to equal* γ_m_ to allow parameter identifiability from protein median levels only. (B) Description of real-time control of gene expression experiments. Every hour, cytometry acquisition is performed, and after gating, deconvolution and FSC-normalization the data is fed to a model-predictive control (MPC) algorithm. The algorithm uses the model to search for the best sequence of duty fractions for 10 duty cycles of period 30 minutes (i.e. a horizon of 5 hours) in order to track the target level. (C) Real-time control results for four different target levels, performed in parallel in different bioreactors (custom setup). Left: median of single cells (controlled value). Right: single-cell distributions over time. Note that a linear scale is used on all plots.

### Exploring the impact of nutrient scarcity on fitness and cellular stress

Fluorescent proteins can be used as reporters to assess phenotypic traits of cells or as barcodes to label strains with specific genotypes^18^. Together with automated cytometry from bioreactor arrays, this capability unlocks a new class of experiments: multiplexed strain characterization and competition in dynamically controlled environments (Figure 4A). Indeed, some fluorescent proteins can be used for genotyping and others for phenotyping. Automated cytometry (including raw data analysis) will then provide quantitative information on both the competition dynamics between the different strains and cell state distribution dynamics for each strain. Depending on the goal of the experiment, this rich information can be fed back to experiment control to adapt environmental parameters for each reactor.

**Figure 4.**
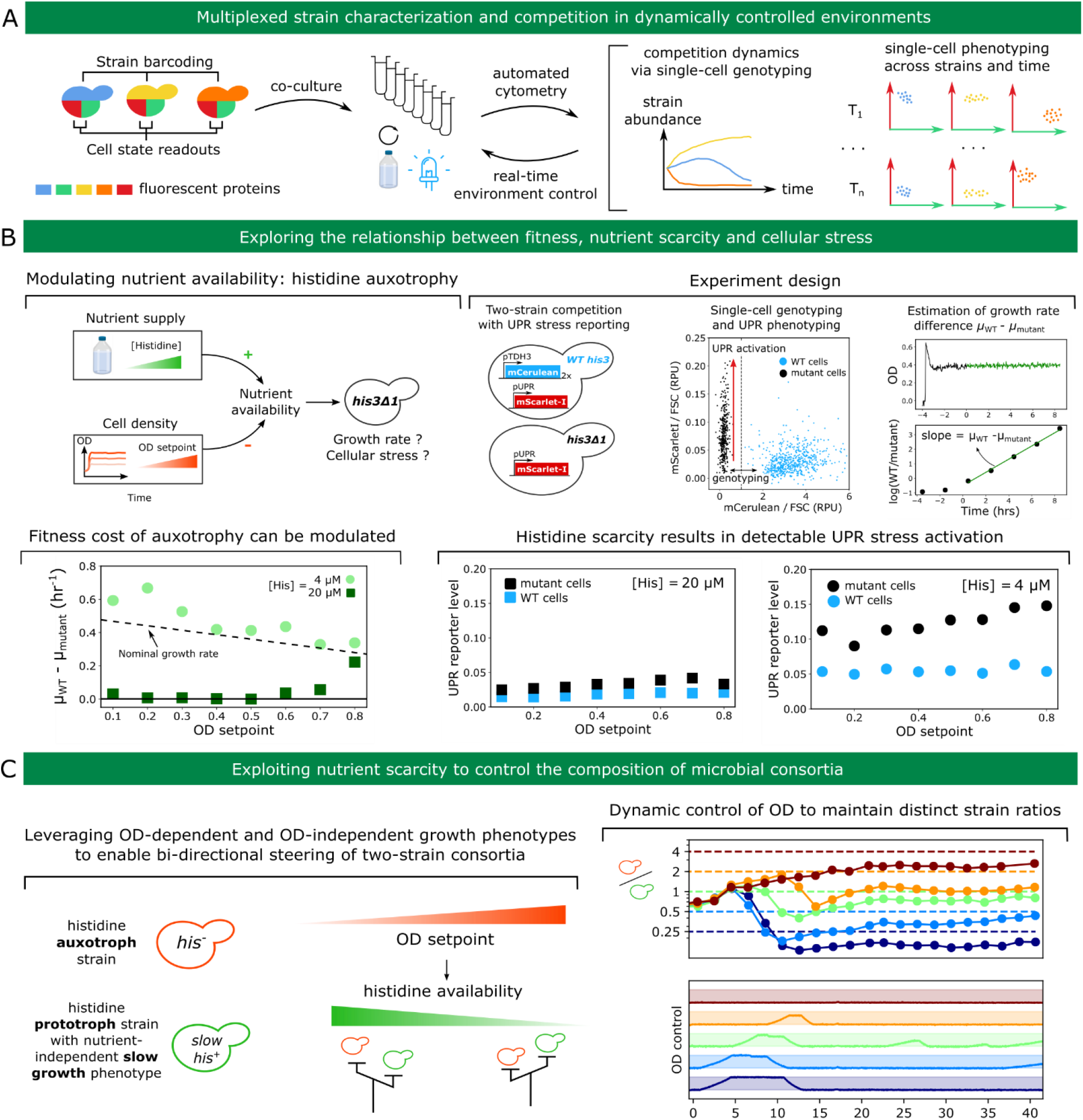
Exploring and exploiting the relationship between fitness, nutrient scarcity and cellular stress. (A) Opening up a new class of experiments by combining co-cultures, automated cytometry for single-cell genotyping and phenotyping and reactive experiment control to adapt environmental conditions in real-time. (B) Top-left: the availability of essential nutrients (such as histidine for his3 mutant strains) depends on the environmental supply but also on cell density via nutrient consumption. Low nutrient availability will impede growth rate and might trigger cellular stress. Top-right: experiment design. Wild-type cells (marked with mCerulean constitutive expression) are co-cultured with his3 mutant cells. Both strains harbor a UPR stress reporter construct driving expression of mScarlet-I. Automated cytometry enables to assign single cells to their genotype and to monitor strain-specific UPR activation. The dynamics of the relative amount of the two strains allows inference of the growth rate difference between mutant and wild-type cells for each condition. Bottom-left: cell density dependence of the fitness deficit of mutant cells at two different media histidine concentration. The dashed line indicates the approximate dependence of wild-type growth rate on the OD setpoint. Bottom-right: strain-specific UPR activation for each condition. (C) Left: principle for a two-strain consortium whose composition can be steered using control of OD. Right: implementation and demonstration. The secretion of a heterologous difficult-to-fold protein is used as a nutrient-independent slow growth phenotype. Dynamic control of the OD setpoint is performed using model-predictive control and the ReacSight event system, similarly to Figure 3B (see Methods). We note the presence of a slight steady-state error, that might originate from a slightly lower long-term growth defect of the secreting strain compared to the estimate based on shorter-term characterization data.

As a first proof of concept that such experiments can be carried out, we set out to explore the impact of nutrient scarcity on fitness and cellular stress (Figure 4B, top-left). Different species in microbial communities have different nutritional needs depending on their metabolic diversity or specialization, and their fitness therefore depends not only on external environmental factors but also on the community itself through nutrient consumption, metabolite release, and other inter-cellular couplings^19,20^. As opposed to competition assays in batch, continuous culture allows to control for such factors. For example, in turbidostat cultures, nutrient availability depends on both nutrient supply (i.e. nutrient levels in the input medium) and nutrient consumption by cells (which primarily depends on the OD setpoint). We used histidine auxotrophy as a model for nutrient scarcity: for his3 mutant cells, histidine is an essential nutrient. By competing his3 mutant cells with wild-type cells at different OD setpoints and different histidine concentrations in the feeding medium, we can measure how nutrient scarcity affects fitness (Figure 4B, top-right). Using a stress reporter in both strains also informs about the relationship between fitness and cellular stress in the context of nutrient scarcity. We focused on the UPR (Unfolded Protein Response^21^) stress response to investigate whether nutrient stress can lead to other, a priori unrelated types of stress, which will be indicative of global couplings in cell physiology.

At a histidine concentration of 4 μM, his3 mutant cells are strongly outcompeted by wild-type cells over the range of OD setpoints (0.1 – 0.8) we considered (Figure 4B, bottom-left). This is not the case anymore at a concentration of 20 μM. At this concentration, the growth rate advantage of wild-type cells is close to zero below an OD setpoint of 0.6 (the remaining histidine is sufficient for his3 mutant cells to grow normally) and becomes larger than 0.2 hr^−1^ at the largest OD setpoint of 0.8 (the remaining histidine is too low and limits growth of his3 mutant cells). Therefore, for this level of nutrient supply, levels of nutrient consumption by cells have a strong impact on fitness of his3 mutant cells. This qualitative change between 4 μM and 20 μM is highly consistent with the reported value of 17 μM for the K_m_constant of the single high-affinity transporter of histidine, HIP1^22^. Also, because the growth rate difference between wild-type and mutant cells for a histidine concentration of 4 μM is close or even exceeds the typically observed growth rate of wild-type cells (between 0.3 and 0.45 hr^−1^ depending on the OD setpoint), we conclude that mutant cells are fully growth-arrested in these conditions. UPR data shows little difference between mutant and wild-type cells across all OD setpoints for a histidine concentration of 20 μM but a clear activation of the UPR response in mutant cells at a histidine concentration of 4 μM (Figure 4B, bottom-right). Therefore, seemingly similar growth phenotypes (such as mutant cells at OD 0.8 for 4 and 20 μM) can correspond to different physiological states (as revealed by differences in UPR activation).

Finally, to showcase reactive control of the environment informed by strain abundance data, we set out to dynamically control the ratio of two strains. Taking control over the composition and heterogeneity of microbial cultures is anticipated to enable more efficient bioprocessing strategies^23^. We reasoned that the OD of the culture could be used as a steering knob when one of the two strain is auxotroph for histidine. Indeed, the strong OD-dependence of the histidine biosynthesis mutant growth rate at a medium histidine concentration of 20 μM (Figure 4B, bottom left) means that switching the OD setpoint of turbidostat cultures can be used to dynamically control its growth rate. In addition, if such strain is co-cultured with a strain prototroph for histidine but growing slower in an OD-independent manner, bi-directional steering of the two strains ratio can be achieved (Figure 4C, left). We built such strain by leveraging burdensome heterologous protein secretion. We then constructed a simple model to predict the (steady-state) growth rate difference with the histidine auxotroph strain (see Methods). Using this model for model-predictive control and the ReacSight event system, we could maintain distinct ratios of the two strains in parallel bioreactors (Figure 4C, right) in a fully automated fashion.

## Discussion

We report the development of ReacSight, a strategy to enhance multi-bioreactor setups with automated measurements and reactive experiment control. ReacSight addresses an unmet need by allowing researchers to combine the recent advances in low-cost, open-hardware instruments for continuous cultures of microbes (e.g. eVOLVER, Chi.Bio^3,4^) and multi-purpose, modular, programmable pipetting robots (e.g. Opentrons OT-2) with sensitive, but generally expensive, stand-alone instruments to build fully automated platforms that open up radically novel experimental capabilities. ReacSight is generic and easy to deploy, and should be broadly useful for the microbial systems biology and synthetic biology communities. While we deployed the ReacSight strategy for only one measurement device (a benchtop cytometer), it should be possible to position two devices on each side of the pipetting robot to enable even more advanced workflows.

As already noted by Wong and colleagues^3^, connecting a multi-bioreactor setup to a cytometer for automated measurements could enable single-cell resolved characterization of microbial cultures across time. Automated cytometry in the context of microbial systems and synthetic biology has in fact already been demonstrated years ago by a small number of labs^6,13,24^, but low throughput or reliance on expensive automation equipment likely prevented a wider adoption of this technology. Automated cytometry from continuous cultures becomes especially powerful in combination with recently developed optogenetic systems^25,26^, enabling targeted, rapid and cost-effective control over cellular processes^13^. We used ReacSight to connect two distinct bioreactor setups (our own, pre-existing custom setup and the recent Chi.Bio^4^ optogenetic-ready bioreactors) with a cytometer. This demonstrate the modularity of the ReacSight strategy, and the platform version using Chi.Bio bioreactors illustrates how other labs lacking pre-existing bioreactor setups could build such platform at a small time and financial cost (excluding the cost of the cytometer, which are expensive but already widespread in labs given their broad usefulness even in absence of automation). We demonstrated the key capabilities of such platform by performing, in a fully automated fashion and in different reactors in parallel, 1) light-driven real-time control of gene expression; 2) cell-state informing competition assays in tightly controlled environmental conditions; and 3) dynamic control of the ratio between two strains.

Still, we only touched the surface of the large space of potential applications offered by such platforms. Strain barcoding can be scaled up to 20 strains with 2 fluorophores and even to 100 strains with 3 fluorophores as recently demonstrated using ribosomal frameshifting^18^. Such multiplexing capabilities can be especially useful to characterize the input-output response of various candidate circuits (or the dependence of circuit behavior across a library of strain backgrounds) in parallel (using different light inductions across reactors). Immuno-beads can be used for more diverse cytometry-based measurements (the robot enabling automated incubation and wash, for example using the Opentrons OT-2 magnetic module). Technologies such as surface display^27,28^ or GPCR signaling^29^ can also be used to engineer biosensor strains to measure even more dimensions of the cultures with a single cytometer and at no reagent costs. Aside of high-performance quantitative strain characterization, such platforms can be useful for biotechnological applications^10^. Automated cytometry informing on the composition of artificial microbial consortia together with dynamic control of culture conditions (as demonstrated here using histidine auxotrophy and OD) could strongly reduce the need to engineer robust coexistence mechanisms^30^, therefore enabling the use of a much larger diversity of consortia.

In the future we hope that many ReacSight-based platforms will be assembled and their design shared by a broad community to drastically expand our experimental capabilities, in order to shed new light on fundamental questions in microbiology or to unlock the potential of synthetic biology in biotechnological applications.

## Methods

### Cloning and strain construction

All integrative plasmids are constructed using the modular cloning framework for yeast synthetic biology Yeast Tool Kit by Lee and colleagues^31^ and all strains originate from the common laboratory strain BY4741. Strain genotypes are described in Table 1 of Text S1.3, and maps of the corresponding integrative plasmids are available [online](https://gitlab.inria.fr/InBio/Public/reacsight/-/tree/master/paper-data/plasmid-maps). All strains used in this work express the light-inducible transcription factor EL222 from the *URA3* locus (transcriptional unit: pTDH3 NLS-VP16-EL222 tSSA1, common parental strain yIB32). Single-color constitutive expression strains (Figure 2) also harbor a pTDH3 FP tTDH1 transcriptional unit at the *LEU2* locus where FP is mCerulean, mNeonGreen or mScarlet-I. Corresponding CDS have been codon-optimized for expression in *S. cerevisiae*. The three-color strain harbors the same three transcriptional units in tandem (order: mCerulean, mNeonGreen, mScarlet-I) at the *LEU2* locus. The autofluorescence strain harbors an empty cassette at the LEU2 locus to match auxotrophy markers between strains. For light-inducible gene expression (Figure 2 and 3), a pEL222 mNeonGreen tTDH1 transcriptional unit (where pEL222 is composed of 5 copies of the EL222 binding site followed by a truncated CYC1 promoter, originally named 5xBS-CYC180pr^16^) is integrated at the *LEU2* locus. For the histidine competition experiments (Figure 4B), the histidine mutant strain (yIB90, parental strain yIB32) expresses a pUPR mScarlet-I tENO1 transcriptional unit integrated at the *LEU2* locus to report on the UPR activation. The pUPR promoter consists in 4 copies of a consensus UPR element^32^ followed by a truncated CYC1 promoter. The histidine wild-type strain was obtained from the mutant strain yIB90 by integrating two identical pTDH3 mCerulean tTDH1 transcriptional units in tandem at the *HO* locus with *HIS3* selection, thereby restoring histidine prototrophy and enabling fluorescent barcoding. For the two-strain consortium experiment (Figure 4C), the slow-growth histidine prototroph strain was obtained by integrating three identical pEL222 alpha-prepro scFv 4-4-20 tTDH1 (burdensome secretion of an anti-fluorescein single chain antibody fragment^33^) transcriptional units in tandem at the *HO* locus (*HIS3* selection) into yIB90 and blue light was used to induce the slow growth phenotype.

### Cell culture conditions

All experiments were performed in 30 mL culture volume bioreactors (cf Text S1.2) at 30 degrees and in turbidostat mode (OD 0.5, typically corresponding to 10^7^ cells/mL according to cytometry data) with synthetic complete medium (ForMedium LoFlo yeast nitrogen base CYN6510 and Formedium complete supplement mixture DCS0019) except for histidine competition experiment where histidine drop-out amino-acid mixture was used (Sigma Y1751) and complemented with desired levels of histidine (Sigma 53319).

### Cytometry acquisition and raw data analysis

Gain settings of our cytometer (*Guava EasyCyte 14HT, Luminex*) for all channels were set once and for all prior to the study such that yeast auto-fluorescence under our typical growth conditions is detectable but at the lower end of the instrument 5-decade range. We verified that cytometry data was reproducible week-to-week with those fixed settings. Single-color strains described above were used together with the autofluorescence control strain to obtain ‘spectral’ signatures of the three fluorophores *mCerulean*, *mNeonGreen* and *mScarlet-I* and autofluorescence levels for each channel. These signatures were also highly reproducible week-to-week (Figure 7A, Text S1.2). To convert raw cytometry data into fluorophore concentrations in relative promoter units (*RPU*^34^), we used a pipeline described in Text S2.2. In essence, it uses data from single-color strains with pTDH3-driven expression for normalization. This pipeline was implemented in *python* (mainly using NumPy^35^ functions) and is available in the *ReacSight* Git [repository](https://gitlab.inria.fr/InBio/Public/reacsight).

### Model-predictive control

For real-time control of gene expression using light (Figure 3), model-predictive control using the two-variables, three-parameters ODE model described in Figure 3A was used. For state estimation upon arrival of cytometry data, the *FP* estimate was set equal to the fluorescence measurement (median of gated, deconvolved data) and the *mRNA* estimate was simply an ‘open-loop’ estimate based on simulating the history of light induction. This first state estimate corresponds to the state of the system at the time of sampling. To account for the time interval (and the concomitant light induction profile) between the sampling time and the data arrival time (typically 10-15 minutes), the model was used to obtain the corresponding updated state estimate. Then, a multi-dimensional, bounded, gradient-based search using SciPy^36^ was used to find the best set of next light duty cycles minimizing the model-predicted distance to the target value over an horizon of 5 hours (10 duty cycles). The corresponding code is available in the *ReacSight* Git [repository](https://gitlab.inria.fr/InBio/Public/reacsight).

### Histidine competition assays

Pre-cultures were performed in synthetic complete medium. Cells were washed in the same low histidine medium as the one used for turbidostat feeding of the competition culture and mixed with an approximate ratio mutant:WT of 5:1 (to ensure good statistics for long enough even when the mutant fitness is very low) before inoculation. Cytometry was acquired automatically every 2 hours. At steady-state, the ratio between two competitors in a co-culture evolves exponentially at a rate equals to their growth rate difference. Linearity of the ratio logarithm for at least 3 timepoints was therefore used to assess when steady-state is reached. A threshold of 1 *mCerulean RPU* was used to assign each cell to its genotype. Size gating was performed as described in Text 2.2 (parameters: size threshold = 0.5 and doublet threshold = 0.5, less stringent than for experiments of Figure 2 and 3) to discard dead or dying cells.

### Dynamic control of the two-strain consortium

A simple sigmoidal model describing the steady-state growth rate difference between the two strains as a function of OD was fitted on previous characterization data corresponding to different OD setpoints. Every two hours, cytometry data was automatically acquired. To assign a genotype to each cytometry event, the combined GRN-B and ORG-G channels was used (the histidine auxotroph strain being GRN-B positive and ORG-G negative). Based on the resulting estimate of the two-strain ratio, the model was used to optimize a vector of future OD setpoints (changing every 2 hours for the next 10 hours) using SciPy^36^.

## Acknowledgments

The authors would like to thanks Cosmin Saveanu, Emmanuel Frachon and Alain Jacquier for the gift of the 16-vessel custom bioreactor setup and for guidance regarding its functioning, enabling us to customize and modernize it even further. We also received guidance and help from Albane Imbert (heading the Institut Pasteur *Fab Lab*) for the custom design and fabrication of Plexiglas and 3D-printed pieces.

## Author contributions

F.B., S.S-C and G.B conceived the study. F.B. performed software and hardware engineering, performed experiments, analyzed data, and developed mathematical models. S.S.-C. developed strains with help of M.F, performed experiments with help of C.A., and analyzed data. A.F. helped with software and hardware developments. F.B. and G.B. supervised the study. F.B., S.S-C and G.B wrote the manuscript with input from all authors.

