## Supplementary Text 1 for "Enhancing bioreactor arrays for automated measurements and reactive control with ReacSight"

#### Enhancing bioreactor arrays for automated measurements and reactive control with ReacSight

*François Bertaux[1,2 *†], Sebastián Sosa-Carrillo[2,1,3 *], Achille Fraisse[2,1], Chetan Aditya[2,1,3], Mariela Furstenheim[1], Gregory Batt[2,1 †]*

1. Institut Pasteur, 28 rue du Docteur-Roux, 75015 Paris, France
2. Inria Paris, 2 rue Simone Iff, 75012 Paris, France
3. Université de Paris, 85 boulevard Saint-Germain, 75006 Paris, France

* These authors contributed to this work equally.

In a first section, we describe in more details the ReacSight strategy, and in a second section, we describe our ReacSight-based platforms enabling reactive optogenetic control and automated cytometry of yeast continuous cultures. Note that two platforms exist, either with a custom bioreactor array or with Chi.Bio bioreactors. In a third section, we provide a detailed description of the strains and plasmids used in this study.

Related code and files are available on the ReacSight Git repository.

### 1. The ReacSight strategy

ReacSight combines both software (section 1.1) and hardware (section 1.2) components.

#### 1.1 Software

ReacSight is based on a versatile multi-instrument control architecture using python and the web-application framework Flask. First, Python is used to write instrument-specific APIs (Figure 1) enabling programmatic control of each instrument (bioreactor components - pumps, optical density readings, LEDs – pipetting robot, measurement device, etc…). As illustrated with three examples, the Python ecosystem of open-source libraries allows to handle a variety of situations.

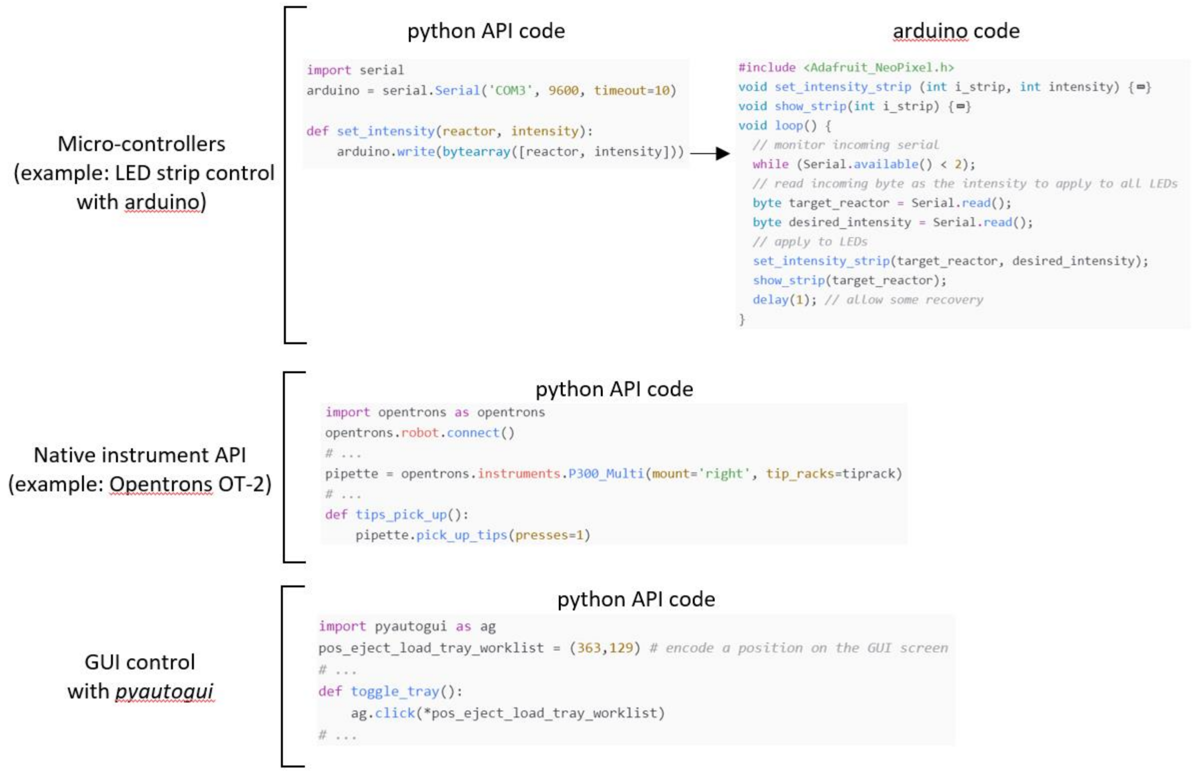


*Figure 1.* ***Programmatic control of various instruments with Python.*** *Code snippets illustrating the versatility of python to control a diversity of instruments using open-source libraries such as pySerial (first example), pyautogui (last example) and opentrons for the Opentrons OT-2 pipetting robot (middle example).*

Next, the web-application framework Flask can be used to expose all instrument APIs to enable control from a single computer (Figure 2). Writing the Flask app code for each instrument is straightforward and requires very little coding. For each instrument, the corresponding app should be running on a computer to which the main experiment control computer has access to. It could be another computer solely dedicated to the control of a given instrument (typically for the measurement device) connected to the local network or the same computer as the one running the experiment (for example for bioreactor components). For the Opentrons OT-2 robot, Flask can be installed and run from the Raspberry Pi computer inside the robot.

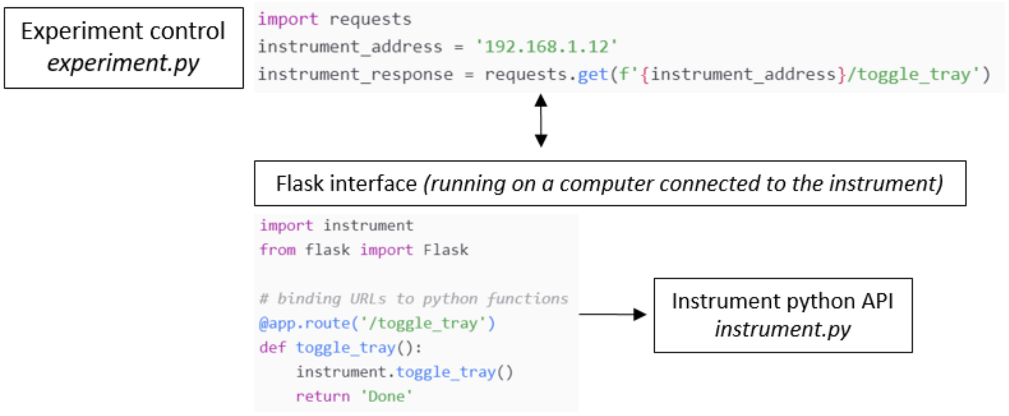


*Figure 2.* ***Using Flask to expose instruments for simple access from a single experiment control computer.*** *Flask enables to develop very easily a web-app interface exposing the instrument control API to any computer who can access the computer running the Flask app (and controlling the instrument). At the level of experiment control, sending commands and receiving the corresponding data is easily done using the module requests. Note that in many situations, the same computer can be used for experiment control and instrument control.*

Finally, once multi-instrument control from a single computer with Python is achieved, the last step to start using a ReacSight platform is to write the code that execute desired experiments. At this stage, several platform-specific and experiment-specific challenges as well as choices to make could arise. Nevertheless, we propose a code architecture (i.e. a recipe for coding platform-specific experiments rather than the code itself) that should cover most scenarios and that should facilitate reactive experiment control and the re-use of code from one experiment to another. The main idea of this code architecture (Figure 3) is to create a clear separation between 1) what is common across experiments and highly constrained by the platform design (main experiment loop) and 2) what can be made highly specific within an experiment and between different reactors in the same experiment (the reactor programs). With this architecture, it becomes natural to write the main experiment loop with a procedural style and the reactor programs with an object-oriented or functional style. To make this more concrete, we provide in the ReacSight Git repository an example implementation of an experiment loop and corresponding reactor programs.

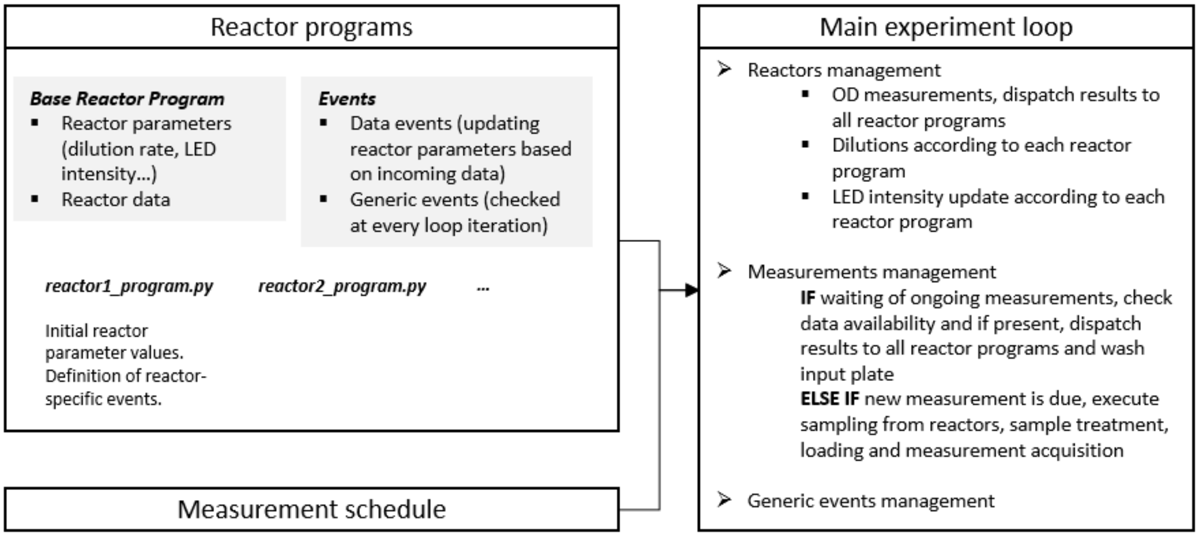


*Figure 3.* ***A python code architecture for reactive experiment control of ReacSight platforms.*** *High-level description of the proposed architecture. The core of the experiment execution will be performed via a single and generic 'main experiment loop' (right). This is where the necessary synchronization for running parallel bioreactor experiments while measuring them using a single robot and measurement device will take place. On the other hand, all the experiment-specific and reactor-specific logic will be described through reactor programs (left). Each reactor program has given parameters than can be dynamically changed during the experiment through the definition of events.*

#### 1.2 Hardware

Core ReacSight hardware components are presented in Figure 4. The two critical elements are the sampling and handling of bioreactor culture samples with the robot (Figure 4A) and the positioning of the measurement device allowing access to the measurement input plate from the robot (Figure 4B). Laser-cut pieces were obtained from 5mm thick acrylic plates cut with a TROTEC Speedy 300 machine ($C0_2$ laser). The convenience (Figure 4C) funnel thrash was 3D-printed with using an Ultimaker 3 machine using Ultimaker Material 2.85 PLA plastic thread. The 3D model was designed in Fusion 360.

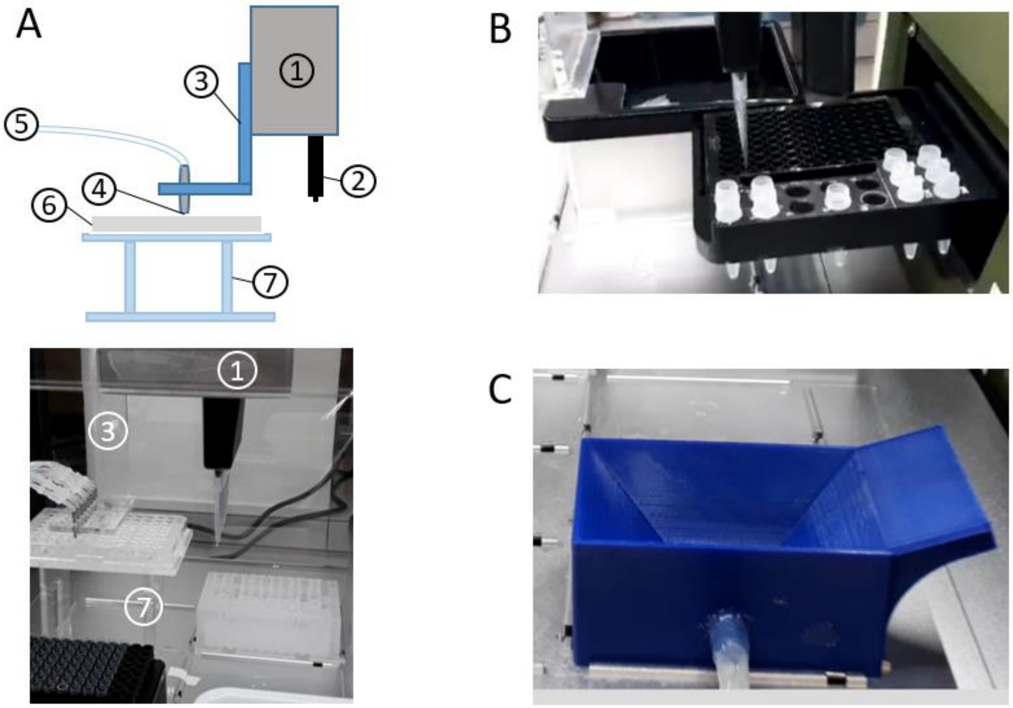


*Figure 4.* ***Core ReacSight hardware components. (A)*** *Sampling and handling of bioreactor culture samples. Top: schematic showing 1 – X-Y moving arm of the robot, 2 – Z moving robot multi-pipette, 3 – sampling head (laser-cut acrylic), 4 – end of bioreactor sampling lines, 5 – sampling lines from bioreactors (should be pump-controlled), 6 – sampling plate, 7 – sampling plate holder (elevation required to prevent the sampling head to collide with other labware elements as the robot arm moves). Bottom: picture of the corresponding setup.* ***(B)*** *Robot access to the instrument input plate for treated samples loading and well washing. Typically, three rows of the input plate are accessible.* ***(C)*** *Convenience 3D-printed (PLA plastic) 'funnel thrash' for disposal of significant liquid quantities.*

#### 2. ReacSight platform instance: reactive optogenetic control and automated cytometry for yeast continuous culture

We provide here details about our platform enabling reactive optogenetic control and automated cytometry for yeast continuous culture.

##### 2.1 Optogenetic-enabled bioreactor setups

We present here the two optogenetic-enabled bioreactor setups used in the two versions of the platform.

###### 2.1.1 A custom 16-reactor platform with optogenetic capabilities

We first extended a custom bioreactor platform designed and built for internal use at the *Institut Pasteur* (E. Frachon, A. Jacquier and C. Saveanu). An overview of the platform is given in Figure 5.

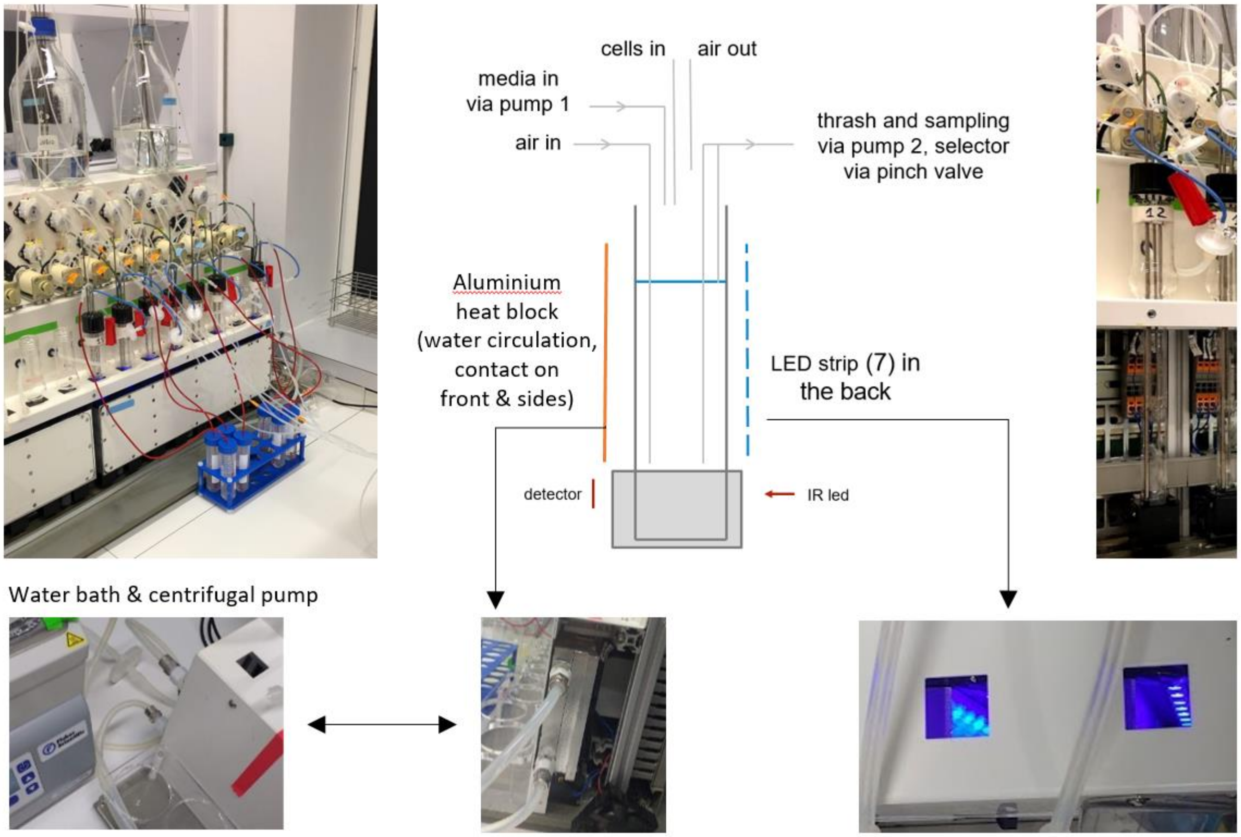


*Figure 5.* ***Overview of the custom-16 reactor platform with optogenetic capabilities used in our lab.*** *Top-left: full view of the bioreactors, showing the vessels, media input bottles, pumps, etc. Top-right: scheme and picture of the vessel design in terms of air and liquid input/outputs. Mixing is achieved thanks to the bubbling caused by the air input. The scheme also highlights how optical density is measured, the position of a LED strip (bottom right picture) on one side of each vessel slot for optogenetic induction, and heating via contact to aluminium blocks on all other sides of the vessel (middle bottom picture). A closed water circulation system from (and back to) a water bath powered by a centrifuge pump maintains the temperature of the aluminium blocks.*

##### 2.1.2 Setup with the Chi.Bio reactors

To demonstrate the modularity of the ReacSight strategy as well as illustrating that a platform enabling reactive optogenetic control and automated cytometry of microbial continuous culture can be easily assembled from low-cost, commercially available optogenetic-ready bioreactors, we constructed an equivalent platform (Figure 6A) based on the Chi.Bio system (Steele et al., PLoS Biology, 2020; purchased from LabMaker, Germany).

Because the native operating system of the Chi.Bio system is already a Flask app, very little changes to the original code (the modified version is available in the ReacSi ght Git repository) were necessary. We validated the functionality and performance of the using the light-induced gene expression strain, obtaining excellent reactor-to-reactor reproducibility of automated cytometry measurements upon a variety of light-induction profiles (Figure 6B).

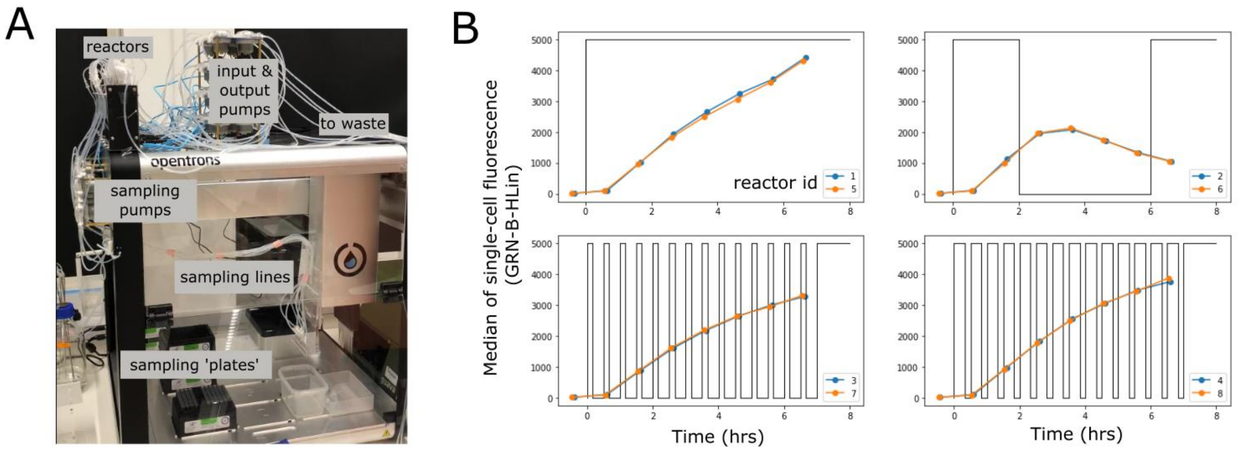


*Figure 6.* ***Description and validation of the platform version using Chi.Bio reactors.*** *(A) Annotated picture of the setup. A few technical differences exist between the two platforms. For example, differences in pumps flow rate resulted in a stronger output stream compared to the other version, requiring us to adapt the type of sampling plates (from regular 300 uL 96-well plate to deeper, larger-volume plates) to eliminate undesired 'splashing'. Also, because with the Chi.Bio setup, three pumps per reactor can be used, a completely separate output line is used for excess volume removal and goes directly to a waste bin. The dead volume is similar for both platforms (around 2 mL). (B) Validation experiments. The eight reactors were run in parallel with the light-induced gene expression strain (same as in Figure 2D and Figure 3 of the main text). Four different ON/OFF blue light induction patterns (dark lines) were applied in pairs of reactors to assess functionality of light-induced gene expression as well as reactor-to-reactor reproducibility. Cytometry data acquisition was performed automatically every hour.*

#### 2.2 Metrology and automated analysis of cytometry data from yeast cultures

We verified that with fixed cytometry acquisition settings as well as fixed turbidostat culture conditions, raw fluorescence distributions of single-color strains are reproducible from week-to-week (Figure 7A). After autofluorescence substraction, each single-color strain gives us a spectral signature across all fluorescence channels of our instrument, and these signatures are also highly reproducible week-to-week and even month-to-month (Figure 7B).

Our automated analysis pipeline to go from raw fluorescence data to size-normalized fluorophore levels (i.e. concentrations for cytosolic fluorophores) is presented in Figure 7C (gating based on forward and side scatter data) and Figure 7D (per-cell fluorescence deconvolution). Note that our gating pipeline also includes a doublet removal step not shown here for simplicity, which excludes cells based on deviation from linearity between the *Area* and *Height* of the forward scatter data. After gating and deconvolution, fluorophore levels are divided by the forward scatter to yield size-normalized fluorophore levels. Finally, to express these values in Relative Promoter Units, they are normalized by equivalent numbers obtained once and for all with single-color strains with pTDH3-driven constructs.

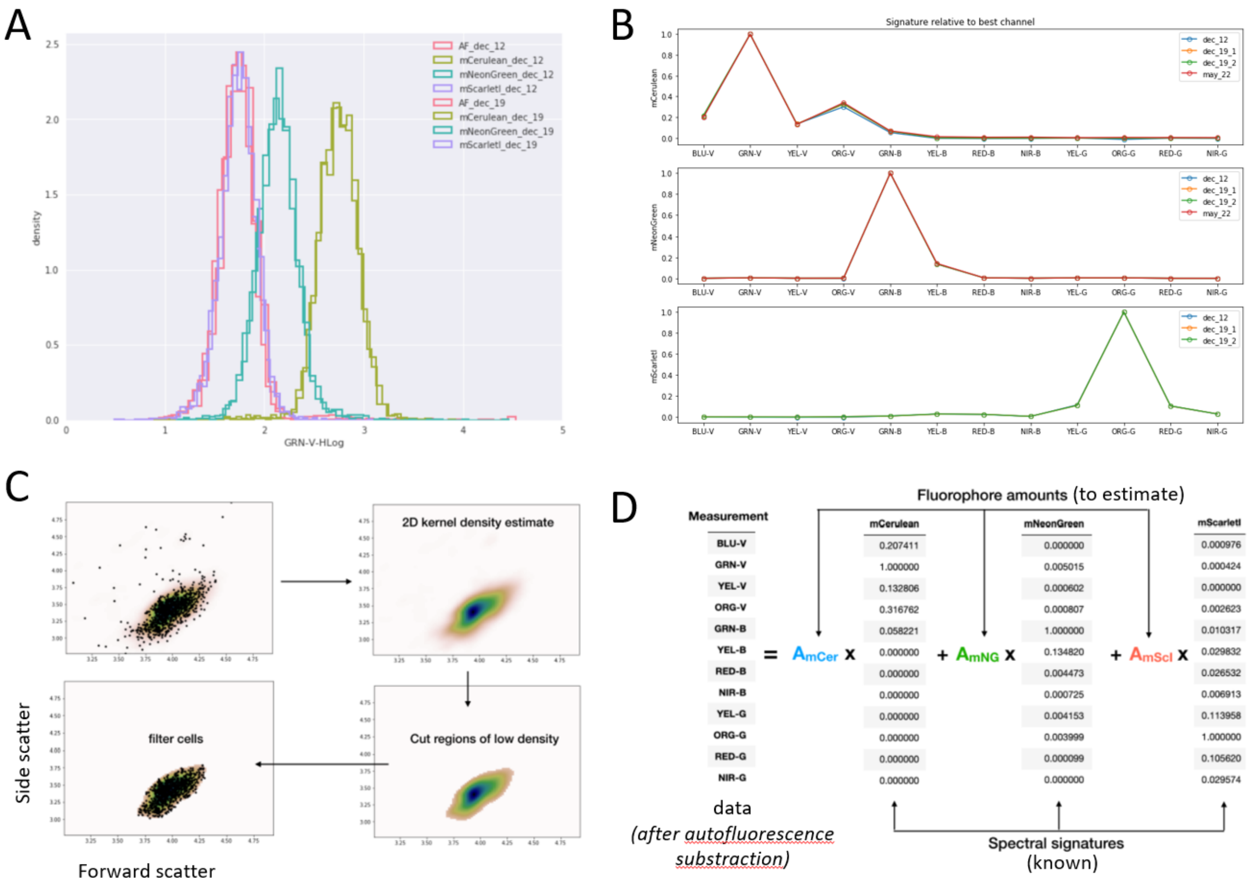


*Figure 7.* ***Metrology and automated analysis of cytometry data from yeast cultures. (A)*** *The same turbidostat culture experiments for single-color strains were performed one week apart and cultures were measured with our cytometer with fixed acquisition settings. Here raw distributions for one channel is shown. Distributions precisely overlapped week-to-week.* ***(B)*** *Spectral signatures for the three fluorophores mCerulean, mNeonGreen and mScarlet-I (i.e., relative channel intensity after autofluorescence substraction for a corresponding single-color strain) are also highly reproducible week-to-week and even month-to-month.* ***(C)*** *Description on the size gating procedure, based on local density in the FSC – SSC space. The doublet removal procedure is not shown here. In total, our gating procedure has two parameters: the size gating threshold (between 0 and 1, cut-off of the local density relative to the maximum density) and the doublet removal threshold (deviation from linearity between FSC Area and Height normalized to the dataset deviation).* ***(D)*** *Per-cell deconvolution is performed by solving the depicted linear system with Numpy linear algebra least-squares solver. Note that the residual error can be used to inform on the quality of the deconvolution on a per-cell basis.*

#### 2.4 Using the OT-2 robot for automated cytometry

The setup enabling automated cytometry with the OT-2 robot is described in more details in Figure 8B. We used fluorescent beads to validate the efficiency of the robot-based washing of the input plate wells inside the cytometer (Figure 8A).

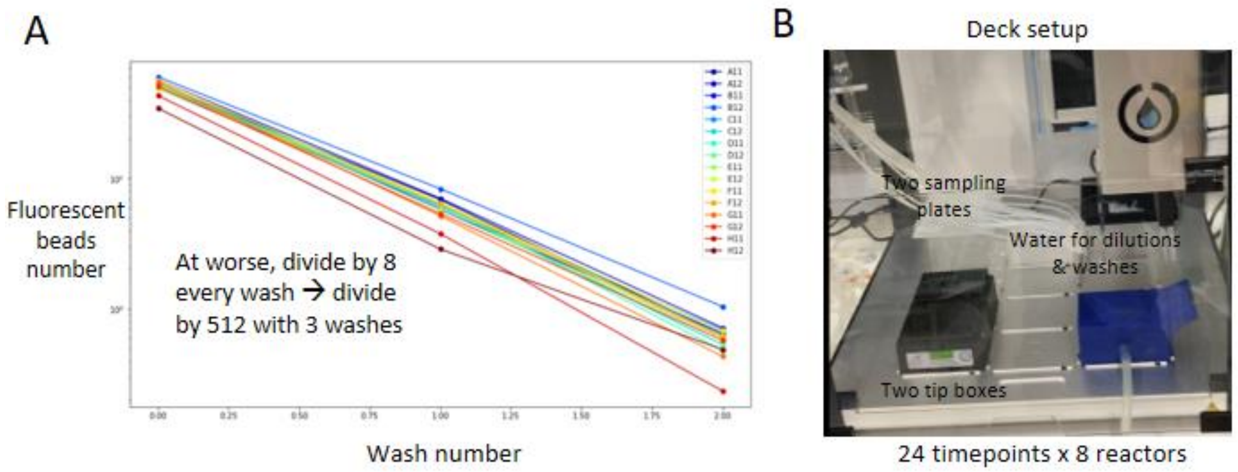


*Figure 8.* ***Cytometer plate washing and deck setup for automated cytometry with the OT-2 robot. (A)*** *Validation of the efficiency of washing steps using fluorescent beads. Consecutive washes from a starting solution containing fluorescent calibration beads are performed and the decrease in beads events per volume is followed. Higher efficiency can be obtained by further minimizing dead volume.* ***(B)*** *Our typical deck setup for starting an experiment. With two sampling plates and two tip boxes, we can perform 24 timepoints for each of 8 reactors in parallel without any human intervention. Renewing tips, plates and water reservoir takes only a few minutes and allow to continue an experiment over several days.*

#### 3. Strains and plasmids used in this study

Table 1 provides a detailed description of all strains described in this study.

| Name | Parent strain | URA3 locus (URA3 selection) | LEU2 locus (LEU2 selection) | HO locus (HIS3 selection) | Auxotrophies | Used in Figure(s) |
|---|---|---|---|---|---|---|
| yIB32 | BY4741 | pTDH3 NLS-VP16-EL222 tSSA1 (pIB120) | | | leucine, histidine, methionine | |
| yIB56 | yIB32 | pTDH3 NLS-VP16-EL222 tSSA1 (pIB120) | spacer (pIB137) | | histidine, methionine | 2B, 2C |
| yIB87 | yIB32 | pTDH3 NLS-VP16-EL222 tSSA1 (pIB120) | pTDH3 mCerulean tTDH1 (pIB273) | | histidine, methionine | 2B, 2C |
| yIB88 | yIB32 | pTDH3 NLS-VP16-EL222 tSSA1 (pIB120) | pTDH3 mNeonGreen tTDH1 (pIB274) | | histidine, methionine | 2B, 2C, 4C |
| yIB43 | yIB32 | pTDH3 NLS-VP16-EL222 tSSA1 (pIB120) | pTDH3 mScarlet-I tTDH1 (pIB135) | | histidine, methionine | 2B, 2C |
| yIB84 | yIB32 | pTDH3 NLS-VP16-EL222 tSSA1 (pIB120) | pTDH3 mCerulean tTDH1 \| pTDH3 mNeonGreen tTDH1 \| pTDH3 mScarlet-I tTDH1 (pIB270) | | histidine, methionine | 2B, 2C |
| yIB86 | yIB32 | pTDH3 NLS-VP16-EL222 tSSA1 (pIB120) | pEL222 mNeonGreen tTDH1 (pIB272) | | histidine, methionine | 2D, 3 |
| yIB90 | yIB32 | pTDH3 NLS-VP16-EL222 tSSA1 (pIB120) | pUPR mScarlet-I tENO1 (pIB115) | | histidine, methionine | 4B |
| yIB119 | yIB90 | pTDH3 NLS-VP16-EL222 tSSA1 (pIB120) | pUPR mScarlet-I tENO1 (pIB115) | pTDH3 mCerulean tTDH1 \| pTDH3 mCerulean tTDH1 (pIB267) | methionine | 4B |
| yIB135 | yIB90 | pTDH3 NLS-VP16-EL222 tSSA1 (pIB120) | pUPR mScarlet-I tENO1 (pIB115) | pEL222 alpha-prepro-4420-scFv tTDH1 \| pEL222 alpha-prepro-4420-scFv tTDH1 \| pEL222 alpha-prepro-4420-scFv tTDH1 (pIB312) | methionine | 4C |

*Table 1.* ***Genotypes of all strains used in this study****. All strains are derived from BY4741 using genomic integration of integrative plasmids constructed with the Yeast Tool Kit modular cloning system (Lee et al., 2015). Names of integrative plasmids are indicated and corresponding plasmid maps (GenBank format) are available online.*
